# Uncharacteristic task-evoked pupillary responses implicate atypical locus coeruleus activity in autism

**DOI:** 10.1101/863928

**Authors:** Michael C. Granovetter, Charlie S. Burlingham, Nicholas M. Blauch, Nancy J. Minshew, David J. Heeger, Marlene Behrmann

## Abstract

Autism spectrum disorder (ASD) is characterized partly by atypical attentional engagement, such as hypersensitivity to environmental stimuli. Attentional engagement is known to be regulated by the locus coeruleus (LC). Moderate baseline LC activity globally dampens neural responsivity and is associated with adaptive deployment and narrowing of attention to task-relevant stimuli. In contrast, increased baseline LC activity enhances neural responsivity across cortex and widening of attention to environmental stimuli regardless of their task relevance. Given attentional atypicalities in ASD, this study is the first to evaluate whether individuals with ASD exhibit a different profile of LC activity compared to typically developing controls under different attentional task demands. Males and females with ASD and age- and gender-matched controls participated in a one-back letter detection test while task-evoked pupillary responses—an established inverse correlate for baseline LC activity—were recorded. Participants completed this task in two conditions, either in the absence or presence of distractor auditory tones. Compared to controls, individuals with ASD evinced atypical pupillary responses in the presence versus absence of distractors. Notably, this atypical pupillary profile was evident despite the fact that both groups exhibited equivalent task performance. Moreover, between-group differences in pupillary responses were observed only in response to task-relevant and not to task-irrelevant stimuli, providing confirmation that the group differences are specifically associated with distinctions in LC activity. These findings suggest that individuals with ASD show atypical modulation of LC activity with changes in attentional demands, offering a possible mechanistic and neurobiological account for attentional atypicalities in ASD.

**Significance Statement:** Individuals with autism spectrum disorder (ASD) exhibit atypical attentional behaviors, such as environmental hypersensitivity and atypical fixedness, but the neural mechanism underlying these behaviors remains elusive. One candidate mechanism is atypical locus coeruleus (LC) activity, as the LC has a critical role in attentional modulation. Elevated LC activity is associated with environmental exploration, while moderate LC activity is associated with focused attention on relevant stimuli. This study shows that, under tightly controlled conditions, task-evoked pupil responses—an LC activity proxy—are lower in individuals with ASD than in controls, but only in the presence of task-irrelevant stimuli. This suggests that individuals with ASD evince atypical modulation of LC activity in accordance with changes in attentional demands, offering a mechanistic account for attentional atypicalities in ASD.

## Introduction

Autism spectrum disorder (ASD) is a neurodevelopmental condition characterized by atypicalities in social, sensory, and motor behaviors, with unclear neural underpinnings (Lord et al., 2018). The diversity of cognitive behaviors implicated in ASD suggests a possible global disruption in the homeostasis of excitatory-inhibitory (E-I) neural activity (Sur and Rubenstein, 2005; Robertson et al., 2013; Dinstein et al., 2015; Rosenberg et al., 2015). Specifically, an inability to modulate neural gain—the likelihood of excitatory versus inhibitory output from a given input (Servan-Schreiber et al., 1990)—could result in increased variability in neural responsivity (Rosenberg et al., 2015). Consistent with this account, functional magnetic resonance imaging (fMRI) studies have demonstrated that individuals with ASD exhibit higher intra-individual variability of stimulus-evoked hemodynamic responses in sensory cortical areas compared to controls (Dinstein et al., 2012; Haigh et al., 2015). This neural variability may be related to or be a product of an inability to regulate neural gain globally.

The locus coeruleus (LC) globally regulates neural gain in association with cognitive task engagement (that is, deployment of attention to task-relevant versus distractor stimuli; Aston-Jones and Cohen, 2005; Eldar et al., 2013). With moderate tonic (baseline) LC activity, phasic responses can be elicited specifically in association with decisions executed on a task, and this mode of activity correlates with increased task engagement. However, with higher tonic LC activity, phasic responses in association with decision processes are weaker, and this mode of activity correlates with decreased task engagement and increased distractibility (Aston-Jones and Cohen, 2005; Gilzenrat et al., 2010). Furthermore, with high tonic LC activity, neural gain is increased throughout cortex, such that neural responsivity is arbitrarily and globally elevated (Aston-Jones and Cohen, 2005).

If individuals with ASD were to exhibit higher tonic LC activity than controls, with consequent increased neural sensitivity throughout cortex (Aston-Jones and Cohen, 2005; Eldar et al., 2013), this might explain the unreliability of neural responses to sensory stimuli in individuals with ASD (Dinstein et al., 2012; Haigh et al., 2015). In fact, individuals with ASD are known to exhibit elevated tonic pupil sizes (Anderson and Colombo, 2009; Anderson et al., 2013; Blaser et al., 2014), and pupil size has been shown to correlate with LC activity in nonhuman primates (Aston-Jones et al., 1994; Joshi et al., 2016). Despite the multiplicity of provocative findings, however, no study has clearly demonstrated whether individuals with ASD evince an atypical LC profile under different attentional demands. A further desideratum of such a study would to be demonstrate differences in LC profiles when behavioral performance is comparable between ASD participants and controls—such an outcome would reveal an inherent alteration in LC activity rather than any physiological differences that might be a direct consequence of differences in behavior.

This study examines whether individuals with ASD exhibit higher tonic LC activity compared to typically developing controls under different attentional demands, by exploiting phasic pupillary responses as a signature of tonic LC activity. The phasic pupillary response to task decisions is an ideal readout of tonic LC activity because it is specifically associated with LC-mediated processing and allows for between-group comparisons that are not confounded by unrelated individual differences in pupil size (Aston-Jones and Cohen, 2005; Eldar et al., 2013). Here, adults with and without ASD performed a one-back letter detection task either in the absence or presence of an auditory distractor. Typically developing individuals, who can flexibly modulate LC activity in the context of attentional demands, are expected to exhibit greater pupillary responses associated with task responses in the presence versus absence of distractors. If, on the other hand, individuals with ASD demonstrate consistently higher tonic LC activity, task-relevant phasic pupillary responses would be expected to be reduced relative to controls’ only in the presence versus absence of distractors and not adapted to the specifics of the task conditions.

## Methods

### Subject Details

Twenty-six individuals with ASD and twenty-six age- and gender-matched controls were initially recruited and participated. The diagnosis of participants with ASD was confirmed by an expert clinician at the Center for Excellence in Autism Research at the University of Pittsburgh, and controls were recruited from the local Pittsburgh community. Descriptive statistics on the Autism Diagnostic Observation Schedule (ADOS) and Wechsler Abbreviated Scale of Intelligence (WASI) for participants with ASD are described in Table 1.

**Table 1.** Clinical metrics of those participants with ASD.

|  | Median | MAD<sup>&</sup> | Minimum | Maximum |
| --- | --- | --- | --- | --- |
| ADOS-LaCo<sup>*</sup> | 4.0 | 1.48 | 2 | 8 |
| ADOS-RSI<sup>^</sup> | 7.0 | 1.48 | 4 | 12 |
| ADOS-LaCo+RSI | 11.0 | 2.97 | 7 | 17 |
| ADOS-RRB<sup>#</sup> | 2.0 | 1.48 | 0 | 6 |
| VIQ<sup>+</sup> | 115.0 | 8.90 | 91 | 141 |
| PIQ<sup>%</sup> | 119.0 | 20.76 | 81 | 134 |
| FSIQ<sup>@</sup> | 113.0 | 19.27 | 86 | 134 |

<sup>\*</sup>LaCo, Language and Communication.
<sup>^</sup>RSI, Reciprocal Social Interaction.
<sup>#</sup>RRB, Restricted and Repetitive Behaviors.
<sup>+</sup>VIQ, Verbal Intelligence Quotient.
<sup>%</sup>PIQ, Performance Intelligence Quotient.
<sup>@</sup>FSIQ, Full Scale IQ.
<sup>&</sup>MAD, median absolute deviation.

Three individuals with ASD and two controls were not included in the data analyses because they did not complete both experimental task conditions (n = 3 participants with ASD, n = 1 control) or because their data was discarded based on artifacts in the data and/or excessive blinks described below (n = 1 control).

In recruitment, groups were matched by age, gender, and handedness (confirmed with the Edinburgh Handedness Inventory; Oldfield, 1971). To determine whether these characteristics were comparable between groups, a logistic regression model to predict group membership was fitted with these features as predictors. Group could not be predicted from a participant’s age (*z* = 1.63, *p* = 0.10), gender (z = 0.05, *p* = 0.96), or handedness (*z* = 0.16, *p* = 0.88), indicating comparability of the groups on these variables. See Tables 2 and 3 for descriptive statistics of these characteristics.

**Table 2.** Age and handedness of participants, by group.

|  | Median |  | MAD |  | Minimum |  | Maximum |  |
| --- | --- | --- | --- | --- | --- | --- | --- | --- |
|  | ASD | Con. | ASD | Con. | ASD | Con. | ASD | Con. |
| Age (yr) | 29.0 | 25.5 | 8.90 | 5.19 | 21 | 21 | 49 | 47 |
| Handedness<br>(EHI#) | 80.00 | 70.00 | 29.65 | 36.19 | -70.00 | -85.71 | 100.00 | 100.00 |

EHI: Edinburgh Handedness Inventory ranges from -100 (left-handed dominance) to +100 (right-handed dominance; Oldfield, 1971). Con., control.

**Table 3.** Percentages of participants (by group) who were female, had consumed caffeine on the day of the study session, were currently taking medications that interact with the adrenergic system, and wore eyeglasses.

|  | ASD | Control |
| --- | --- | --- |
| Female | 8.70% | 8.33% |
| Caffeine | 43.48% | 54.17% |
| Adrenergic Medication(s)* | 56.52% | 4.17% |
| Eyeglasses | 52.17% | 41.67% |

\*, significant predictor of group.

Participants completed questions about additional variables that might affect pupillometry measurements. As caffeine intake can affect pupil size (Abokyi et al., 2017), participants were asked about their caffeine intake on the same day of the study session. Participants also listed the medications they were taking, and the UpToDate database (Wolters Kluwer) was used to determine which, if any, medications interact with the adrenergic system. Finally, whether a participant was wearing eyeglasses was noted as this could potentially affect pupillometry recordings. A logistic regression model to predict group was fitted with these features as predictors. Group membership was predicted by use of adrenergic-related medication (*z* = 3.16, *p* < 0.01), but not by caffeine intake (*z* = 1.37, *p* = 0.17) or wearing eyeglasses (*z* = 0.16, *p* = 0.87; Table 3). The effect of medication use was thus accounted for in the analyses described below.

The Carnegie Mellon University Institutional Review Board reviewed and approved this research, and all participants provided informed consent.

### Experimental Design and Statistical Analyses

#### Task Design

Participants’ heads were positioned in a chinrest at a distance of approximately 60 cm from an approximately 38-by-31 cm computer monitor. The luminance and contrast settings of the monitor, as well as the ambient lighting in the room, were approximately constant throughout the experimental session and across participants. Task stimuli were presented using the Psychophysics Toolbox (Brainard, 1997) in MATLAB (MathWorks), and participants completed two versions of the task: without and with accompanying distractors.

The luminance of all stimuli was comparable to the background: specifically, the L* value of the CIELAB color space (McGuire, 1992) was approximately equal for all colors in the task display. On a gray (CIELAB = [5776.9 0 0]) background, participants viewed a green (CIELAB = [5777 -4812.8 4645.1]) circle positioned at the center of the screen. A set of 15 lower-case gray (CIELAB = [5776.9 0 0]) letters randomly appeared one at a time at a frequency of 2 Hz within the circle. Participants were instructed to indicate, using a keyboard press, each instance in which a consecutive letter repeat occurred. To provide feedback to participants, when a key was pressed in the second following a consecutive letter repetition, the letter on display became purple (CIELAB = [5772.4 3020.8 -5570]) for a 0.5 s duration subsequent to the key press. For all other key presses, the letters became red (CIELAB = [5780.3 5857.9 5501.7]) for a 0.5 s duration subsequent to the key press (Figure 1). While letter presentation was random, a pair of consecutive letters would not repeat within a 6-s interval. The letters were presented in the same order to all participants, and out of a total of approximately 1584 letter presentations, there were 54 total consecutive letter repetitions. This constituted the no-distractor condition.

**Figure 1.**
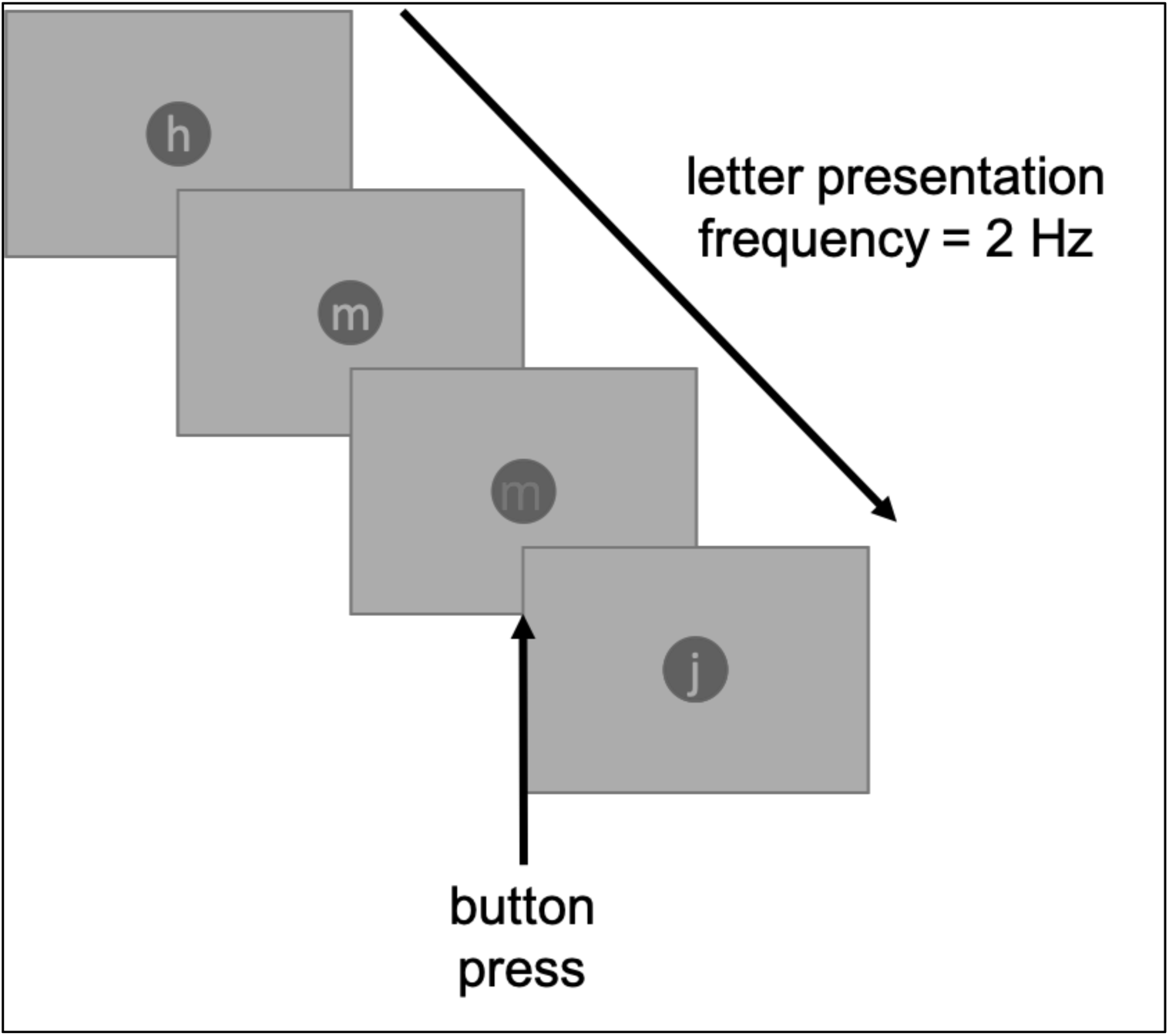
Schematic of the one-back letter detection task. Participants viewed individual presentations of letters at a rate of 2 Hz and were instructed to press a button upon observing a consecutive letter repeat. For a duration of 0.5 s, letters became purple or red in response to a correct or incorrect button press, respectively. The visual display was isoluminant throughout the task session. In the first half of the experiment, participants performed the task in the absence of distractor stimuli. In the second half, participants were exposed to series of tones played temporally independent of the task sequence.

Participants then completed the same task, but this time in the presence of distractor auditory stimuli, following a task design adapted from prior studies (Dinstein et al., 2012; Haigh et al., 2015). As the participants performed the same letter-repeat task, 11 600-Hz tones were played through a headset, each tone lasting for 0.15 s, with 0.15-s intervals between tones. Initiation of the 11 tones was separated by a random intertrial interval, ranging between 6-10 s to prevent participants from predicting the onset of the tones, and the timing of tone onsets was not associated with letter presentations or repeats. During this block, a total of approximately 1607 letters were presented, with 65 total consecutive letter repetitions.

Each of the two task conditions consisted of 3 blocks of letter presentations, with breaks in-between.

#### Eye-tracking

Pupil area and coordinates were measured with the EyeLink 1000 (SR Research, Ottawa, Canada; http://www.sr-research.com/) at a sampling rate of 1000 Hz. The eye-tracker was positioned below the computer monitor and was angled to record measurements from a single eye. A 3- or 5-point display grid was used for calibration, conducted prior to each experimental block. Thresholds for pupil detection were adapted for each participant due to individual differences between participants, such as participants’ needs to wear glasses or contact lenses, eye color, and eye size. To determine if these parameters of the eye-tracker were comparable between the groups, a logistic regression model to predict group was fitted with the thresholds for pupil and cornea detection as predictors. Neither pupil detection threshold (ASD: median = 80.00, median absolute deviation (MAD) = 7.41; Control: median = 80.00, MAD = 7.41; *z* = 1.47, *p* = 0.14) nor cornea detection threshold (ASD: median = 250.00, MAD = 0; Control: median = 250.00, MAD = 0; *z* = 0.01 *p* = 0.99) was predictive of group.

The pupillometry data were preprocessed using custom in-house scripts in MATLAB version 9.5.0 (MathWorks), as well as adapted blink/artifact interpolation code (Urai et al., 2017). Pupil area was converted to pupil diameter, taking into account the fact that the eye-tracker used a centroid-fitting model in detecting the pupil. Instances in which the eye-tracker could not track the pupil, and instances in which the pupil size was beyond three standard deviations (SD) from the median pupil size of the block were considered to be artifacts. During blinks and artifacts (including those detected by the EyeLink 1000 software), the data were linearly interpolated over these intervals and nearest neighbor interpolation was used at the start and end points of these intervals. Blinks, partial blinks, or other artifacts detected within 0.25 s of one another were linearly interpolated as a single blink, and data were linearly interpolated from 0.15 s prior to and 0.15 s after each detected blink. Nearest neighbor interpolation was employed at the start and end of each blink/artifact. To interpolate over peak-detected blinks, the pupil size data were initially smoothed using a two-dimensional digital filter with an 11-point symmetric Hann window. Peak-detected blinks (separated in time by a minimum duration of 0.5 s) were subsequently interpolated: peak-detected blinks detected within 0.25 s of one another were interpolated as a single peak-detected blink, and data were interpolated from 0.3 s prior to and 0.15 s subsequent to each peak-detected blink. Nearest neighbor interpolation was also employed at the start and end of each peak-detected blink. Furthermore, to meet criteria for inclusion in the study, a participant’s data were excluded if blinks or artifacts constituted more than two-thirds of the duration of an experimental condition (absence versus presence of distractors) across all blocks for that condition. (Only one participant, a control, did not meet this criterion, and his data are not included in the summary statistics above nor in the analyses below.)

To assess each participant’s baseline pupil size, at the start of each block, participants viewed a central fixation (the same green circle on a gray background used in the experiment) for approximately 45 s prior to starting the task. For each participant, the median of all pupil size measurements across these passive viewing periods was computed. One participant (in the ASD group) blinked and exhibited artifacts for more than two-thirds of the duration of baseline pupil size recordings. This participant’s data were thus discarded from analyses of baseline pupil size only.

Parameters for preprocessing of the pupillometry data were decided upon prior to completion of data collection and final performance of statistical analyses, based on visual inspection of initial data collection. For analyses of task-evoked pupil responses, pupil size measurements were converted to percent signal change relative to the mean pupil size within the entire block in which they were collected. This was done to normalize between-block differences in pupil response amplitudes caused by interaction between the tonic and phasic components of the pupil signal (Eldar et al., 2013). To eliminate very low frequency fluctuations, the pupil size signal was high-pass filtered with a Butterworth filter of order 4 with a cutoff of 0.03 Hz. To reduce the sampling rate of the signal for further analysis, a low-pass Chebyshev Type I filter was used with an order of 8, and the sampling rate of the data was subsequently reduced by a factor of 25.

Linear deconvolution was used to estimate how the pupil responded to task events. Deconvolution analysis is a form of regression often used in fMRI analyses where physiological responses to fast stimulus presentations from each trial can introduce noise into the signal for an event of interest (Glover, 1999; McCloy et al., 2016). To “deconvolve” an impulse response function (IRF) of the pupillary response to a given event, the pupil time series data is multiplied by the pseudoinverse of the design matrix with the events of interest (Gardner et al., 2008). For each participant, the pupil’s IRF was deconvolved to a letter repeat preceding a hit, a letter repeat preceding a miss, and the 1-s preceding a false alarm (FA), separately for each task condition (no distractor vs. distractor). A single deconvolution block matrix was used, composed of 3 concatenated design matrices, one per event type, to covary out the other predictors in each IRF’s estimation. It was assumed that each IRF was 4 s in duration. The amplitude of the pupil response was calculated as the median absolute deviation (MAD) of the IRF. If a given pupil response amplitude value was greater than or less than 3 SD from the mean of the pupil amplitudes of all participants in a group (by diagnosis) for an event (hit, FA, or miss), that value was assumed to be artifactual, treated as an outlier, and discarded. Additionally, in a separate analysis, a deconvolution block matrix was used, with a single design matrix with the onset of distractor tones, to generate IRFs (also 4 s in duration) for pupillary responses to distractors.

#### Inferential Analyses

All inferential statistics were performed with R version 3.5.2 (R Foundations for Statistical Computing), using the dplyr (Wickham et al., 2019), psych (Revelle, 2019), lme4 (Bates et al., 2019 p.4), lmerTest (Kuznetsova et al., 2019), and multcomp (Hothorn et al., 2019) packages. Analysis figures were generated using the seaborn Python package (Michael Waskom et al., 2018).

For analyses on sensitivity index (d’), criterion (C), reaction time (RT), and pupil amplitude response (each to hits, FAs, and misses), linear mixed models were fitted to predict these variables, with group and task condition as fixed effect predictors and participant as a random effect predictor. For analyses on average baseline pupil size (for which there is only one derived measurement per participant), linear models were fitted to predict these variables, with group as a predictor. Because use of adrenergic-related medications was predictive of group, this variable was also included as a predictor in all of the aforementioned models. For each dependent measure, separate models were fitted, either including or excluding the use of medications as a predictor, and the Bayesian information criterion (BIC) was calculated for each to determine the optimal model. For models that included medication use as a predictor, the BIC was higher than the model with this variable excluded (the model with the lowest BIC is preferred; Wagenmakers, 2007; Table 4), in all cases. Thus, reported models only include group and task condition (when applicable) as predictors of the respective dependent measure of interest.

**Table 4.**
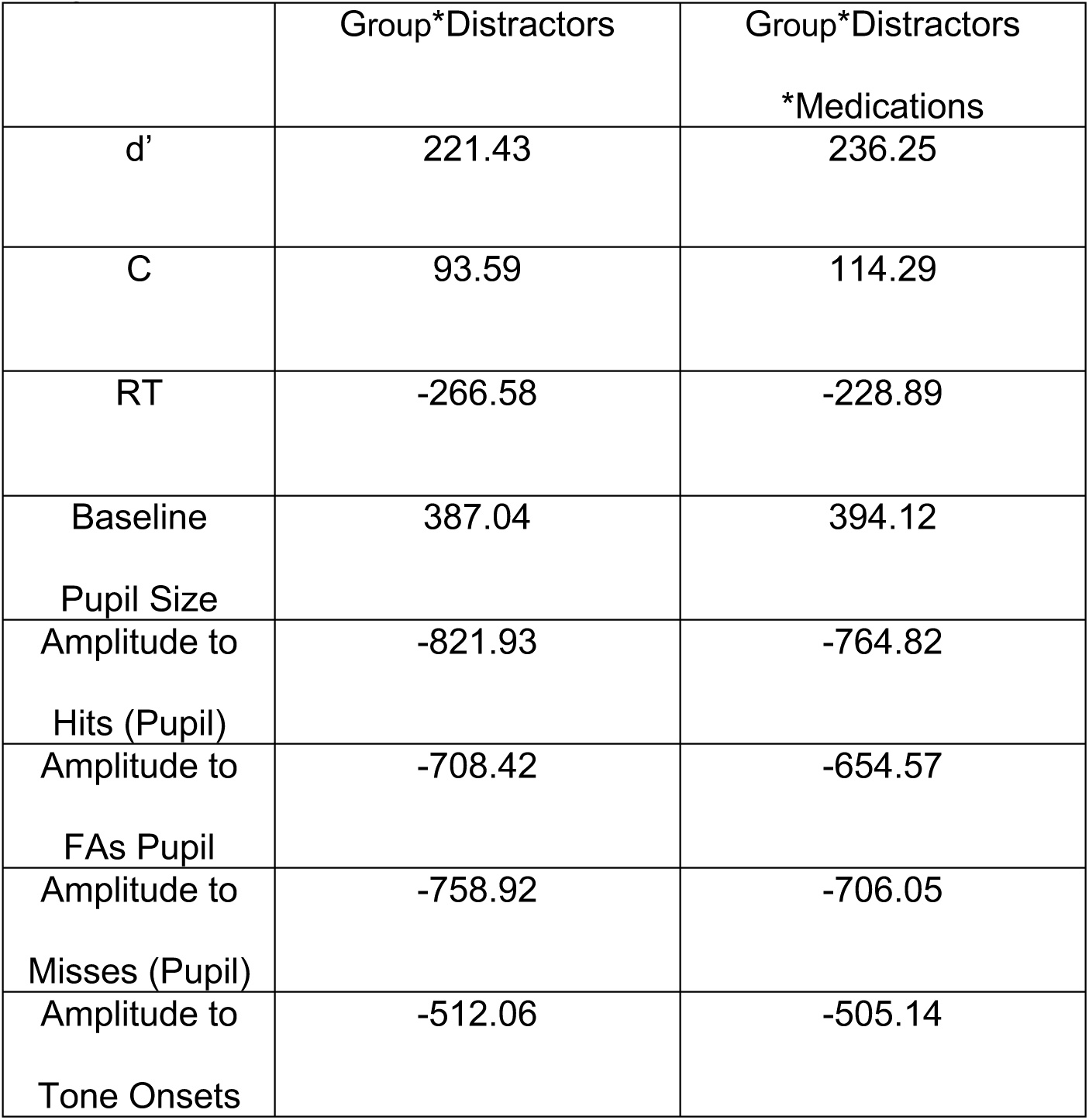
BIC values for models with and without medication use as a predictor. Each cell designates a different model: the row designates the dependent measurement and the column designates the predictors included.

In addition, to verify the findings from the linear mixed models predicting pupil response amplitude to hits, FAs, and misses, for each such task event, the ratio of the pupil response amplitude in the presence of distractors to that in the absence of distractors was computed for each participant. For each task event, a linear model was fitted with this ratio as the dependent measure and group as the predictor.

Absolute values of test statistics are reported. The *α* criterion for statistical significance was designated as 0.05 for all inferential statistical analyses. In cases in which there was no statistical significance, an approximation for the Bayes Factor (BF) was computed using the respective BICs of the null model (excluding all fixed effect predictors) and alternative model (including all fixed effect predictors). A BF between 3 and 20, between 20 and 150, or greater than 150 was designated as positive, strong, or very strong evidence for the null hypothesis, respectively (Wagenmakers, 2007). All participants whose data were not determined to be outliers as described above were included (n = 23 and 24 in the ASD and control groups, respectively). Some participants do not have select data values (such as a participant who does not commit any FAs, and therefore has no pupil amplitude response to FAs); degrees of freedom (df), however, are reported for all inferential analyses.

#### Classification Analyses

To validate the inferential statistical analyses, classification analyses were used to assess whether group membership could be predicted from pupil response amplitude. A logistic regression model was fitted with group as the dependent measure and the absolute difference of the pupil response amplitude between the two conditions (absence versus presence of distractors) as the predictor, for each event type (hit, FA, or miss). The LogisticRegression class within the scikit-learn version 19.1 (Pedregosa et al., 2011) package in Python version 3.7.1 (Python Software Foundation) was used with the saga solver and no regularization. Twenty repeats of five-fold cross-validations were performed to compute the predictive accuracy of group for each event and condition combination. A null distribution was created by shuffling the labels 10,000 times and performing the same cross-validation classification approach. The statistical significance (*p*-value) of the classification accuracy was determined by a comparison to the null distribution, as (1-percentile), where percentile indexes the percentile of the true classification accuracy in the distribution of null distribution classification accuracies.

As three independent classification analyses were performed (3 event types), for these analyses, an accuracy value was considered significant if the *p*-value was lower than the Bonferroni-corrected criterion: 0.05/3 = 0.02.

### Code Accessibility

Experiment and preprocessing MATLAB scripts, R and Python analysis code, and preprocessed data are available on GitHub: https://github.com/michaelgrano/ASD_nback.

## Results

First, group differences in behavioral performance were analyzed to determine whether both groups performed comparably on the task. Second, group differences in time-averaged pupil size were analyzed to rule out the possibility of any systematic a priori differences in pupil size between the groups. Last, between-group comparisons of the pupil response amplitude to each task event (hits, FAs, and misses) for each task condition (absence versus presence of distractors) were analyzed. Group differences in pupil amplitude to distractor tone onsets were also assessed. Pupil amplitude in response to stimuli that elicit hits and FAs, but not to stimuli that elicit misses or to distractor stimuli themselves, are “task-evoked” and should be associated with LC activity because only pupillary responses to cognitive decisions can be inferred to be caused by fluctuations in LC activity (Aston-Jones and Cohen, 2005). Finally, classification analyses were used to determine whether a diagnosis of ASD could be predicted from task-evoked pupil responses alone.

### Comparable between-group task performance in the absence and presence of distractor stimuli

Group differences in behavioral performance were initially analyzed as any such differences could confound observed group differences in pupillary responses. The d’, C, and RT were computed for each participant, for each task condition (absence versus presence of distractors). (If a participant’s d’ or C was positive or negative infinity, the maximum or minimum value for that participant’s group in the given condition was substituted for these analyses, respectively.) Figure 2 shows the d’, C, and RT for the two groups. There was no significant effect of group (*t*(56.29) = 1.57, *p* = 0.12), task condition (*t*(45.00) = 0.66, *p* = 0.52), or their interaction (t(45.00) = 0.16, *p* = 0.88) on d’ (Figure 2a). Likewise, there was no significant effect of group (*t*(66.31) = 1.02, *p* = 0.31), task condition (*t*(45.00) = 1.86, *p* = 0.07), or their interaction (*t*(45.00) = 0.49, *p* = 0.62) on C (Figure 2b). There was very strong evidence that neither group nor presence of distractor stimuli predicts d’ (BF = 3262.08) or C (BF = 8760.19).

**Figure 2.**
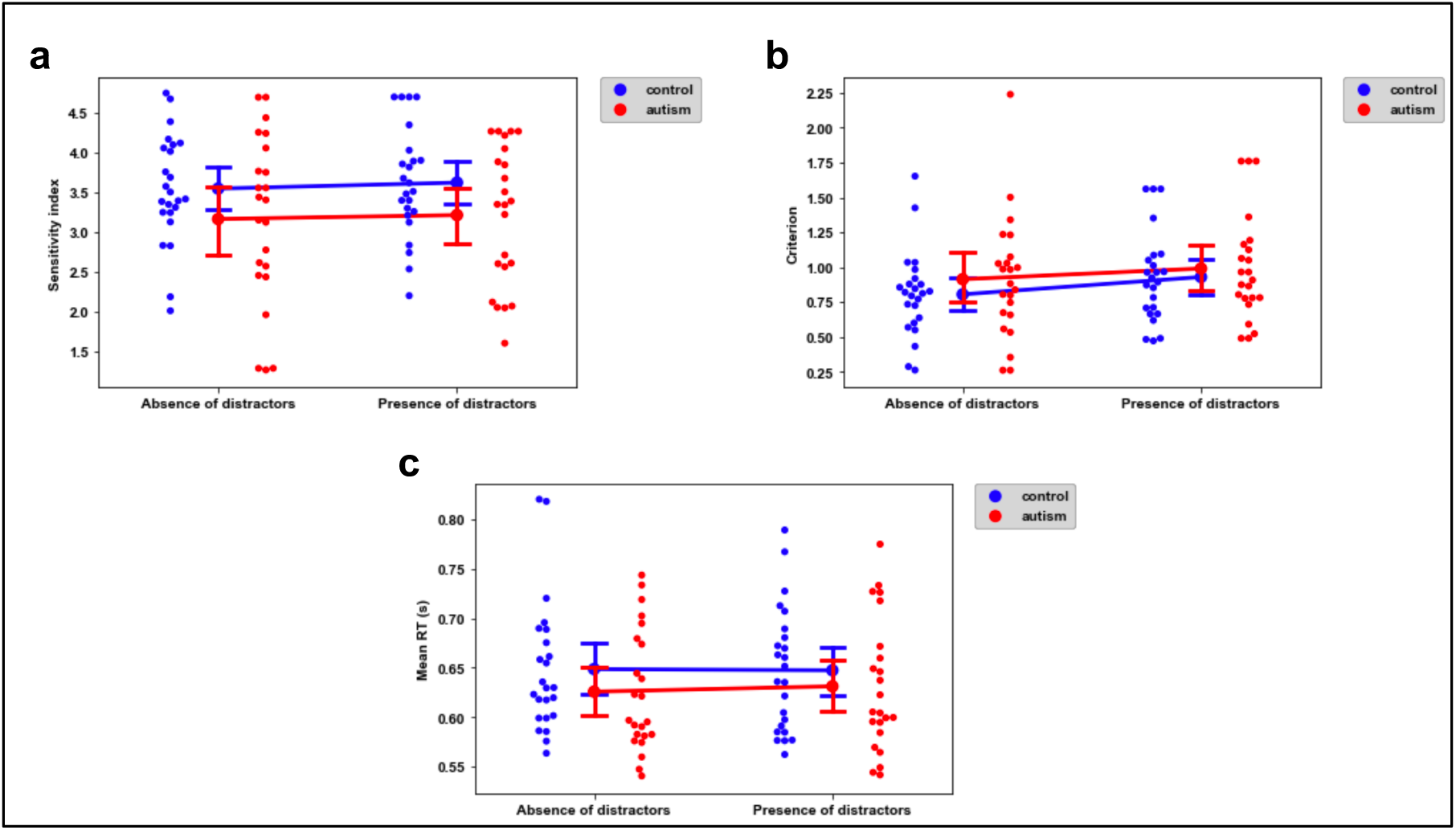
Behavioral performance on the letter detection task. **a**, d’, **b**, C, and **c**, RT, across group (autism versus control) and condition (absence versus presence of distractor stimuli). Each point represents an individual participant. Line plots show mean ± one standard error of the mean (SEM).

The mean RT (time between the onset of a letter repeat and a correct button press) across all correct responses was also computed for each participant, separately for each task condition. There was no significant effect of group (*t*(50.47) = 1.21, *p* = 0.23) or task condition (*t*(45.00) = 0.19, *p* = 0.85) on mean RT. There was also no significant interaction of group × task condition on mean RT (*t*(45.00) = 0.73, *p* = 0.47), and there was very strong evidence in favor of the null hypothesis (BF = 30545766.18; Figure 2c).

The lack of a main effect of group on d’, C, or RT indicates similarity in task performance between the two groups. Given that there are no differences in performance, any differences in pupil size are unlikely to be attributed to differences in behavioral performance and, indeed, a simple task was selected specifically to equate performance as much as possible. The interaction between group × condition also rules out a foundational difference in working memory, a required component of the task, in the ASD versus control participants.

### No between-group differences in baseline pupil size

Group differences in time-averaged pupil size were analyzed to rule out the possibility of any systematic a priori differences in pupil size between the groups. Baseline pupil size (recorded prior to each task block) was compared between groups to determine whether pupil size differed between participants with ASD and controls, independent of the letter detection task. As shown in Figure 3, there was no significant effect of group on the median baseline pupil size (*t*(44) = 0.09, *p* = 0.93), with positive evidence that group does not predict this measure (BF = 6.75). The lack of a main effect of group indicates that there were no systematic differences in pupil size, thereby ruling out confounding variables that would be independent of the task.

**Figure 3.**
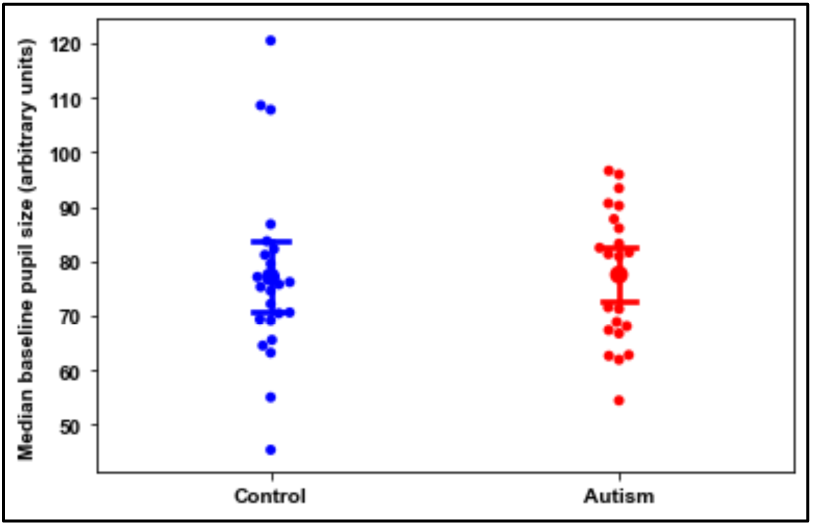
Median baseline pupil size of participants, by group. Each point represents an individual participant. Line plots show mean ± one SEM.

### Individuals with ASD exhibited smaller task-evoked pupil response amplitudes than did controls in the presence but not absence of distractor stimuli

Linear deconvolution analysis (Glover, 1999; McCloy et al., 2016) was used to approximate a 4-s IRF to each task event (hits, FAs, and misses) for each participant in each task condition. The individual IRFs of participants’ pupillary responses to hits are shown in Figure 4. The pupil response amplitude was calculated as the MAD of the IRF, as this value captures the dispersion of the pupillary response, while reducing the impact of noise caused by limited data (Kret and Sjak-Shie, 2019). This is similar to the approach extensively adopted in the fMRI literature, where the dispersion of the blood oxygen level dependent signal time course has been used as a non-parametric measure of response amplitude (Power et al., 2018).

**Figure 4.**
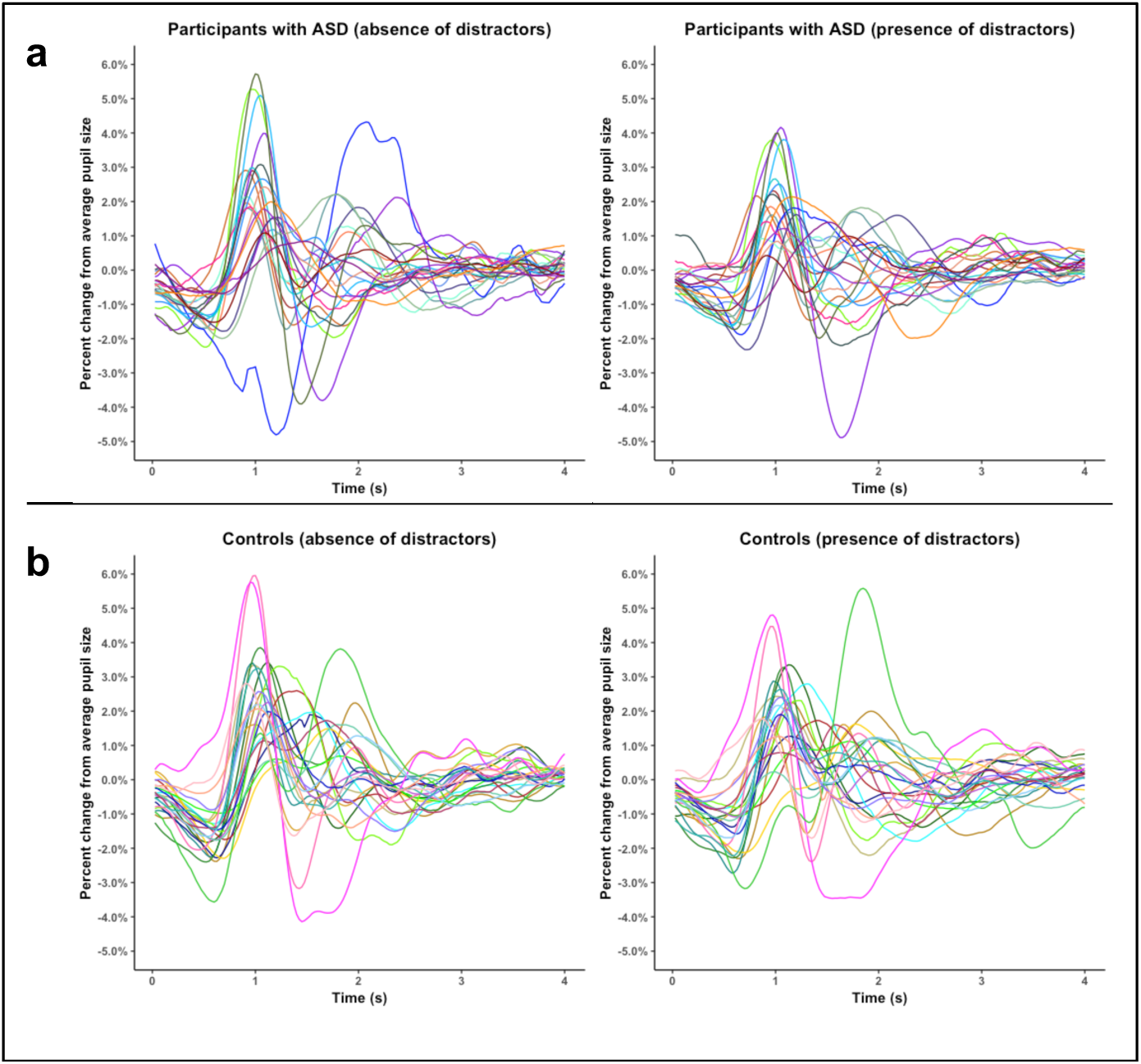
All participants’ individual IRFs of pupillary responses to hits. Each color represents the IRF of a unique participant in the **a**, ASD and **b**, control groups, with the left panel showing the results in absence of distractors and the right panel showing the results in the presence of distractors.

Between-group comparisons of the pupil amplitude response to each task event associated with a decision (hits and FAs) for each task condition (absence versus presence of distractors) were analyzed. These pupillary responses should reflect changes in LC activity because pupil dilations occur specifically in response to the appearance of a stimulus on a cognitive task (here, the one-back letter detection task) that results in a decision (here, a key press; Aston-Jones and Cohen, 2005).

As evident from Figure 5, there was a significant interaction between group and the presence/absence of distractor stimuli on pupil amplitude in response to both hits (*t*(43.63) = 3.06, *p* < 0.01) and FAs (*t*(42.44) = 2.65, *p* = 0.01). Furthermore, in the presence versus absence of distractor stimuli, there was a significant increase in pupil amplitude in response to hits (*t*(43.28) = 2.93, *p* < 0.01), independent of group, but no significant difference in response to FAs (*t*(42.78) = 1.13, *p* = 0.26). Moreover, there was no significant effect of group on pupil amplitude in response to either hits (*t*(67.36) = 0.08, *p* = 0.94) or FAs (*t*(70.46) = 0.17, *p* = 0.87).

**Figure 5.**
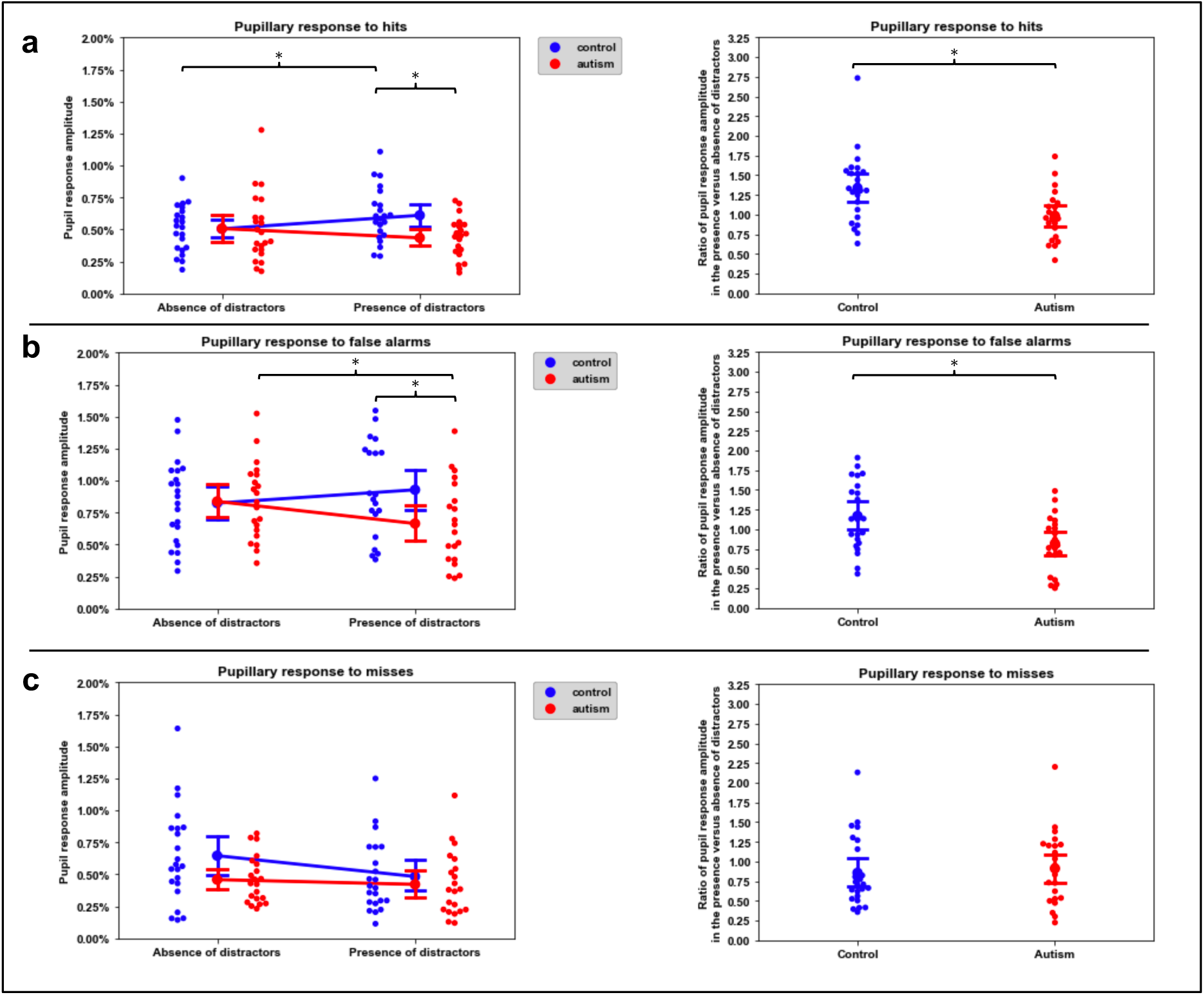
Pupil response amplitudes to ***a***, hits ***b***, false alarms, and ***c***, misses, compared across groups and task conditions. Left-hand panels show pupil response amplitude, as defined as the MAD of the IRF of the respective pupil response, after normalization of the pupil time series data to the mean pupil size of the respective experiment block. Right-hand panels show the ratio of the pupil response amplitude in the presence of distractors to that in the absence of distractors. Line plots show mean ± one SEM. * and ** signify *p* < 0.05 and 0.01, respectively for contrast tests.

Post-hoc contrast tests of the effect of task condition on pupil response amplitude performed separately for each group showed that, as anticipated (Gilzenrat et al., 2010), among controls, the pupil amplitude in response to hits was significantly higher in the presence versus absence of distractors (*z* = 2.93, *p* < 0.01). Notably, there was no such significant difference in response to hits among participants with ASD (*z* = 1.43, *p* = 0.15). Moreover, while there was no significant between-group difference in pupil amplitude in response to hits in the absence of distractors (*z* = 0.08, *p* = 0.94), in the presence of distractors, individuals with ASD exhibited lower pupil response amplitudes than did the controls (*z* = 2.80, *p* < 0.01). In fact, the ratio of the pupil response amplitude in the presence of distractors to that in the absence of distractors was significantly higher among controls than it was among participants with ASD (*t*(45.00) = 3.10, *p* < 0.01; Figure 5a).

Furthermore, among individuals with ASD, the pupil amplitude in response to FAs was significantly lower in the presence versus absence of distractors (*z* = 2.61, *p* < 0.01), while this was not the case among controls (*z* = 1.13, *p* = 0.26). As was the case with hits, the pupil amplitudes in response to FAs were not different between groups in the absence of distractors (*z* = 0.17, *p* = 0.87), but, in the presence of distractors, individuals with ASD exhibited lower pupil response amplitudes than controls (*z* = 2.57, *p* = 0.01). Additionally, the ratio of the pupil response amplitude in the presence of distractors to that in the absence of distractors was significantly higher among controls than it was among participants with ASD (*t*(42.00) = 2.85, *p* = 0.01; Figure 5b).

Thus, overall, pupillary responses to stimuli that elicit behavioral reports (thereby suggestive of LC activity; Aston-Jones and Cohen, 2005) in the distractor-present condition (i.e. with increased cognitive load and task engagement) were lower among individuals with ASD. However, in the absence of distractor stimuli, no between-group differences existed.

### No interaction effect between group and task condition on pupil amplitude responses to misses

If the interaction effect between group and task condition on pupil response amplitude was specific to cognitive effort on the task (which would implicate LC activity (Aston-Jones and Cohen, 2005), then we would not expect to see an interaction of group and task condition on pupil amplitude in response to misses (that is, on trials where effort was likely to be least). Indeed, there was no significant interaction between group and task condition on pupil amplitude in response to misses (*t*(44.00) = 1.41, *p* = 0.17). Additionally, the ratio of the pupil response amplitude in the presence of distractors to that in their absence was not significantly different between the two groups (*t*(45.00)= 0.41, *p* = 0.68; BF = 6.21, positive evidence for the null hypothesis). However, there were main effects of group and of task condition. Individuals with ASD exhibited lower pupil amplitudes in response to misses, independent of task condition, relative to controls (*t*(70.72) = 2.28, *p* = 0.03), and participants in both groups exhibited lower pupil amplitudes in response to misses in the presence versus absence of distractors (*t*(44.00) = 2.92, *p* < 0.01; Figure 5c). It is conceivable that controls might notice misses across both task conditions more so than individuals with ASD, which might explain why controls’ pupil amplitudes in response to misses are overall higher.

In summary, in response to an event that is likely to be only weakly implicated with LC activity because of the lack of cognitive effort to a miss (Aston-Jones and Cohen, 2005), individuals with ASD do exhibit lower pupil response amplitudes, but importantly, independent of the attentional demands of the task.

### Group membership can be predicted from the difference in pupil amplitude responses in the presence versus absence of distractors, only during hits and FAs (and not misses)

To validate the differential response to hits and FAs for the two groups and the effect of distractor condition, an assumption-free classification algorithm was used to determine whether group could be predicted from the task-evoked pupil response amplitudes alone. For each event type (hits, FAs, and misses), a logistic regression model was fitted to assess whether group classification (autism or control) could be predicted from the difference in the pupil response amplitude between the two conditions (absence versus presence of distractors). Consistent with the findings demonstrating an interaction between group and task condition on pupil response amplitudes to hits and FAs, group could be decoded from the between-conditions difference in pupil amplitude in response to hits (accuracy = 0.63, *p* < 0.01) and FAs (accuracy = 0.56, *p* < 0.01) with above-chance accuracy. At the same time, the between-conditions difference in pupil amplitude in response to misses was not predictive of group and was below-chance in accuracy (accuracy = 0.38, *p* = 1.00).

### No between-group differences of pupil dilations to the task-irrelevant distractor stimuli

Group differences in pupil response amplitude just to distractor tone onsets were also assessed. While pupil dilations can occur in response to auditory stimuli (Zekveld et al., 2018), LC activity is not associated with pupil dilations to task-irrelevant stimuli (Aston-Jones et al., 1999; Aston-Jones and Cohen, 2005; Gilzenrat et al., 2010) such as the onset of orthogonal distractors. If differences in pupillary dynamics between the two groups is specific to inherent group differences in LC activity, differences in pupil dilations to task distractor stimuli would not be expected. To test this, pupil amplitude responses to the onset of distractor stimulus presentation were compared between the two groups. There was no significant effect of group on pupil response amplitude to the onset of distractors (*t*(44) = 1.66, *p* = 0.10), suggesting that group does not predict pupillary response to distractors per se (BF = 1.68; Figure 6). Thus, group differences in pupillary dynamics are likely to be independent of pupil responses to the distractor stimulus presentations themselves.

**Figure 6.**
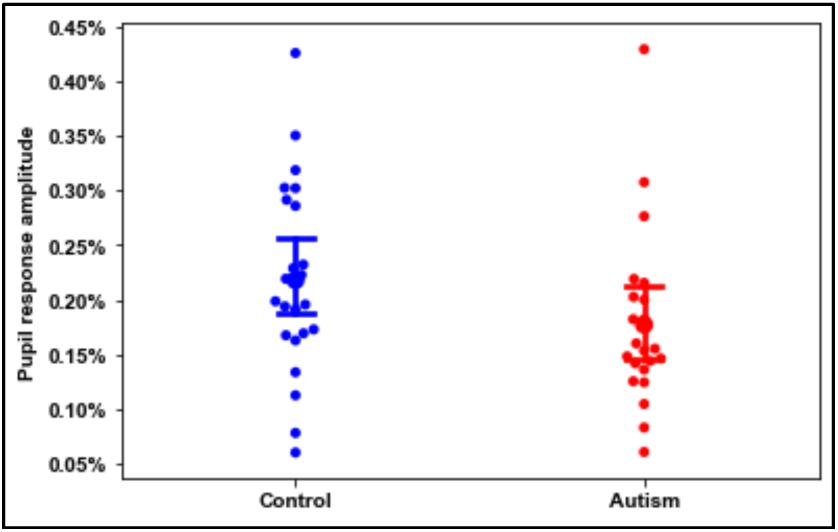
Pupil response amplitude to distractor stimuli, after normalization of the pupil time series data to the mean pupil size of the respective experiment block. Line plots show mean ± one SEM.

## Discussion

The goal of this study was to explore differences in LC activity (inferred from pupillometry measurements) between individuals with ASD and matched controls as they performed a simple visual working memory task in the absence or presence of distractor auditory tones. The ASD and control groups performed the task with statistically indistinguishable accuracy and speed, both in the presence and absence of distractors. However, specifically in the presence of distractors, individuals with ASD exhibited lower task-evoked pupil response amplitudes than did controls. Furthermore, group could be decoded with above-chance accuracy based solely on the difference of task-evoked pupil response amplitudes in the presence versus absence of distractors. This physiological effect could not be accounted for due to between-group differences in medication use or baseline pupil size and was specific to task-evoked responses. As LC activity can be inferred from pupillary responses to task-relevant information, the lower task-evoked pupil response amplitudes in the ASD compared to control participants in the presence of distractors implicates dysregulation of LC activity.

Pupil dilation—specifically in association with a task decision—is an established direct correlate of phasic LC activity and an established inverse correlate of tonic LC activity. A rise in tonic LC activity increases the gain of neural activity indiscriminately throughout cortex, thereby increasing neural responsivity. It has been posited that this indiscriminate increase in cortical gain allows for increased behavioral flexibility, exploration of the task environment, and thus attention to both task-relevant and task-irrelevant stimuli. In contrast, when tonic LC activity and global gain are reduced, attentional deployment shifts to task-relevant stimuli, and attention to task-irrelevant distractors becomes attenuated (Aston-Jones et al., 1999; Aston-Jones and Cohen, 2005; Gilzenrat et al., 2010; Pfeffer et al., 2017). It is thus particularly notable that in the present study participants with ASD evinced lower pupil response amplitudes in the *presence* of distractors because one would expect a typically developing individual to demonstrate increased pupil response amplitude (indicating higher LC activity) under increased attentional demands (Aston-Jones and Cohen, 2005; Gilzenrat et al., 2010).

A recent review from Bast et al. (2018) suggests LC dysfunction might be associated with attentional differences in ASD, but there has been little prior empirical evidence to support this hypothesis. Several studies have shown differences in phasic pupillary responses in ASD (Martineau et al., 2011; Blaser et al., 2014; Nuske et al., 2014a, 2014b; Krach et al., 2015; Lawson et al., 2017), some even suggesting, in contrast to the results here, that individuals with ASD exhibit larger phasic pupillary responses compared to typically developing controls (Blaser et al., 2014; Lawson et al., 2017; Bast et al., 2019). However, in these studies participants with ASD exhibited differences in task performance compared to controls, and thus differences in pupil dilations may be attributable to differences in task performance. Importantly, in the current study, participants with ASD and controls showed comparable performance on the task, in terms of both signal detection theoretic and RT measures. A one-back task was specifically utilized because: 1) it was expected to elicit phasic pupillary responses associated with task decisions, and 2) it was expected that participants with ASD would perform comparably to controls (Williams et al., 2005). Had participants with ASD performed more poorly than controls, the observed interaction of group and task condition on pupil response amplitudes might have been a consequence of task performance rather than of LC activity per se. However, as task performance did not differ between the two groups, the between-group differences in task-evoked pupil response amplitudes across conditions suggest an inherent difference in LC physiology among the participants with ASD.

A number of processes could account for differences in pupillary dynamics in individuals with ASD, and it has been suggested that individuals with ASD exhibit heightened autonomic activity (Cheshire, 2012; Kushki et al., 2013) potentially in relation to comorbid anxiety diagnoses (Lord et al., 2018). Furthermore, prior literature has suggested that individuals with ASD exhibit larger tonic pupil sizes (Anderson and Colombo, 2009; Anderson et al., 2013; Blaser et al., 2014). However, unlike prior findings that might be attributed to generalized autonomic arousal, the results of the present study indicate that differences in pupillary dynamics in the participants with ASD are specifically task-dependent, and, therefore, provide clear inference of LC activity per se. First, group differences were noted, even after controlling for the potential contribution of caffeine or adrenergic-related medications and even though time-averaged pupil size recorded prior to each task block was equivalent between groups. Second, an interaction of task condition and group on pupil response amplitudes was only revealed for phasic pupillary responses in association with hits and FAs, but not in association with misses. Third, there were no group differences in phasic pupil amplitudes in response to distractor stimulus onset, which alone would not be expected to specifically elicit LC activity (Aston-Jones and Cohen, 2005). These are critical observations because LC activity has been correlated primarily with task-related decisions (Aston-Jones et al., 1999; Aston-Jones and Cohen, 2005; Gilzenrat et al., 2010). Thus, the effects in pupillary dynamics uncovered in this investigation are most likely to be associated with group differences in LC activity.

Elevated tonic LC activity can globally increase neural gain, amplifying neural activity and enhancing overall neural responsiveness (Aston-Jones and Cohen, 2005). Generally, enhanced neural gain is advantageous as it allows for exploratory behaviors and learning from new features of one’s environment. Studies in typically developing individuals suggest that, when in a tonic LC mode, individuals are more likely to attend selectively to salient stimulus cues (Eldar et al., 2016) or focus their attention on stimulus features to which they are individually predisposed to attend to (Eldar et al., 2013). However, an inability to regulate neural gain could limit the ability to distinguish relevant versus irrelevant stimuli (Gilzenrat et al., 2010), thereby hampering the establishment of priors and the ability to learn from novel environmental input (Sinha et al., 2014; Dinstein et al., 2015). Thus, if individuals with ASD exhibit consistently elevated gain, this could enhance attention to particular environmental stimuli, but would impair the ability to properly establish priors (Sinha et al., 2014). Attention might thus be deployed indiscriminately to task-relevant and task-irrelevant stimuli. This indiscriminate but selective attention from elevated gain could explain the fixated interests, selective attention, and hypersensitivity to environmental stimuli in individuals with ASD (Remington et al., 2009; Lord et al., 2018), as consistently high LC tonic activity would ultimately preclude the diversion of attention from distractor or task-irrelevant features in one’s environment (Gilzenrat et al., 2010).

If an individual with ASD cannot readily increase gain in the presence of distractors, this would significantly hamper typical learning processes. Social communication atypicalities in individuals with ASD might also be explained by consistently elevated gain: an inability to learn from social cues and expressions (Lord et al., 2018) might be a reflection of a broader inability to learn from environmental stimuli (Sinha et al., 2014).

If individuals with ASD exhibit higher tonic LC activity than controls in an environment with both task-relevant and irrelevant stimuli, such a dysregulation of the LC system would be consistent with the proposal of disrupted E-I homeostasis of cortical activity in ASD (Sur and Rubenstein, 2005; Rosenberg et al., 2015). Much research on E-I homeostasis has focused on the roles of glutamate and GABA in achieving this balance (Hensch, 2005; Samardzic et al., 2018), which is critical for efficient perceptual processing (Zhou and Yu, 2018). In fact, there have been several demonstrations of atypical GABA activity in ASD (Pizzarelli and Cherubini, 2011; Robertson et al., 2016; Uzunova et al., 2016; Ajram et al., 2017). But perhaps the disruption in E-I homeostasis is not only or strictly a disruption in the *ratio* of excitatory to inhibitory activity, but in the *gain*, which is a measure of the simultaneous amplification (or dampening) of excitatory and inhibitory activity (Servan-Schreiber et al., 1990; Aston-Jones and Cohen, 2005; Hoshino, 2005; Pfeffer et al., 2017). Unregulated neural responsivity due to elevated LC activity would, in fact, be consistent with findings that uncover highly variable neural responses to sensory stimuli in ASD (Dinstein et al., 2012; Haigh et al., 2015). In other words, with dysregulated gain in ASD from elevated LC activity, neural output would be highly unpredictable from neural input.

Consistently elevated tonic LC activity—and consequently globally increased cortical gain—in an attention-demanding environment is thus consistent with clinical and behavioral characteristics of ASD, as well as the E-I homeostasis disruption hypothesis. This study provides physiological evidence for an inherent difference in regulation of tonic LC activity on a task on which individuals with ASD perform comparably to controls, laying the foundation for future work to explore the direct effects of this dysregulation on more challenging tasks that reflect the burdensome cognitive load of one’s real-world environment. These results provide novel evidence for the LC’s role in gain dysregulation, and consequent atypical attention, in ASD and, in addition, offer a possible neurobiological basis for some signatures of ASD such as social communication and restricted learning.

## Acknowledgements

This research was supported by Award Numbers T32GM008208 and T32GM081760 from the National Institute of General Medical Sciences to MCG and a grant from the Simons Foundation Autism Research Initiative to MB and DH. The content is solely the responsibility of the authors and does not necessarily represent the official view of the National Institute of General Medical Sciences or the National Institutes of Health. The authors thank Drs. Yael Niv and Eran Eldar for their initial discussions that stimulated the early framework of this study. The authors also thank Dr. Shaun E. Eack, Patricia J. McCarroll, and Michelle Perrin for assisting with recruitment and John J. Markiewicz for his organization of the neuropsychological testing data. The authors also thank Drs. Ilan Dinstein and Eran Eldar for input on the study design, and Drs. Ilan Dinstein and Sarah M. Haigh, whose experiment code was adapted for the study task. The authors also thank Dr. Timothy Verstynen and Madhumita Harish for input on the analyses. Finally, the authors thank the participants for making this research possible.

## References

Abokyi S, Owusu-Mensah J, Osei KA (2017) Caffeine intake is associated with pupil dilation and enhanced accommodation. Eye 31:615–619.

Ajram LA, Horder J, Mendez MA, Galanopoulos A, Brennan LP, Wichers RH, Robertson DM, Murphy CM, Zinkstok J, Ivin G, Heasman M, Meek D, Tricklebank MD, Barker GJ, Lythgoe DJ, Edden RAE, Williams SC, Murphy DGM, McAlonan GM (2017) Shifting brain inhibitory balance and connectivity of the prefrontal cortex of adults with autism spectrum disorder. Transl Psychiatry 7:e1137.

Allen G, Courchesne E (2001) Attention function and dysfunction in autism. Front Biosci J Virtual Libr 6:NaN–NaN.

Anderson CJ, Colombo J (2009) Larger tonic pupil size in young children with autism spectrum disorder. Dev Psychobiol 51:207–211.

Anderson CJ, Colombo J, Unruh KE (2013) Pupil and salivary indicators of autonomic dysfunction in autism spectrum disorder. Dev Psychobiol 55:465–482.

Aston-Jones G, Cohen JD (2005) AN INTEGRATIVE THEORY OF LOCUS COERULEUS-NOREPINEPHRINE FUNCTION: Adaptive Gain and Optimal Performance. Annu Rev Neurosci 28:403–450.

Aston-Jones G, Rajkowski J, Cohen J (1999) Role of locus coeruleus in attention and behavioral flexibility. Biol Psychiatry 46:1309–1320.

Aston-Jones G, Rajkowski J, Kubiak P, Alexinsky T (1994) Locus coeruleus neurons in monkey are selectively activated by attended cues in a vigilance task. J Neurosci Off J Soc Neurosci 14:4467–4480.

Bast N, Banaschewski T, Dziobek I, Brandeis D, Poustka L, Freitag CM (2019) Pupil Dilation Progression Modulates Aberrant Social Cognition in Autism Spectrum Disorder. Autism Res 0 Available at: https://onlinelibrary.wiley.com/doi/full/10.1002/aur.2178 [Accessed November 5, 2019].

Bates D, Maechler M, Bolker B, Walker S, Christensen RHB, Singmann H, Dai B, Scheipl F, Grothendieck G, Green P, Fox J (2019) lme4: Linear Mixed-Effects Models using “Eigen” and S4. Available at: https://CRAN.R-project.org/package=lme4 [Accessed May 25, 2019].

Blaser E, Eglington L, Carter AS, Kaldy Z (2014) Pupillometry Reveals a Mechanism for the Autism Spectrum Disorder (ASD) Advantage in Visual Tasks. Sci Rep 4:4301.

Brainard DH (1997) The Psychophysics Toolbox. Spat Vis 10:433–436.

Caplan B, Mendoza JE (2011) Edinburgh Handedness Inventory. In: Encyclopedia of Clinical Neuropsychology (Kreutzer JS, DeLuca J, Caplan B, eds), pp 928–928. New York, NY: Springer New York. Available at: https://doi.org/10.1007/978-0-387-79948-3_684 [Accessed May 18, 2019].

Cheshire WP (2012) Highlights in clinical autonomic neuroscience: New insights into autonomic dysfunction in autism. Auton Neurosci 171:4–7.

Dinstein I, Heeger DJ, Behrmann M (2015) Neural variability: friend or foe? Trends Cogn Sci 19:322–328.

Dinstein I, Heeger DJ, Lorenzi L, Minshew NJ, Malach R, Behrmann M (2012) Unreliable Evoked Responses in Autism. Neuron 75:981–991.

Eldar E, Cohen JD, Niv Y (2013) The effects of neural gain on attention and learning. Nat Neurosci 16:1146–1153.

Eldar E, Niv Y, Cohen JD (2016) Do You See the Forest or the Tree? Neural Gain and Breadth Versus Focus in Perceptual Processing. Psychol Sci 27:1632–1643.

Gardner JL, Merriam EP, Movshon JA, Heeger DJ (2008) Maps of Visual Space in Human Occipital Cortex Are Retinotopic, Not Spatiotopic. J Neurosci 28:3988–3999.

Gilzenrat MS, Nieuwenhuis S, Jepma M, Cohen JD (2010) Pupil diameter tracks changes in control state predicted by the adaptive gain theory of locus coeruleus function. Cogn Affect Behav Neurosci 10:252–269.

Glover GH (1999) Deconvolution of Impulse Response in Event-Related BOLD fMRI1. NeuroImage 9:416–429.

Haigh SM, Heeger DJ, Dinstein I, Minshew N, Behrmann M (2015) Cortical variability in the sensory-evoked response in autism. J Autism Dev Disord 45:1176–1190.

Hensch TK (2005) Critical period plasticity in local cortical circuits. Nat Rev Neurosci 6:877–888.

Hoshino O (2005) Cortical Modulation of Synaptic Efficacies through Norepinephrine. In: Adaptive and Natural Computing Algorithms (Ribeiro B, Albrecht RF, Dobnikar A, Pearson DW, Steele NC, eds), pp 70–73. Vienna: Springer.

Hothorn T, Bretz F, Westfall P, Heiberger RM, Schuetzenmeister A, Scheibe S (2019) multcomp: Simultaneous Inference in General Parametric Models. Available at: https://CRAN.R-project.org/package=multcomp [Accessed May 25, 2019].

Joshi S, Li Y, Kalwani RM, Gold JI (2016) Relationships between Pupil Diameter and Neuronal Activity in the Locus Coeruleus, Colliculi, and Cingulate Cortex. Neuron 89:221–234.

Krach S, Kamp-Becker I, Einhäuser W, Sommer J, Frässle S, Jansen A, Rademacher L, Müller-Pinzler L, Gazzola V, Paulus FM (2015) Evidence from pupillometry and fMRI indicates reduced neural response during vicarious social pain but not physical pain in autism. Hum Brain Mapp 36:4730–4744.

Kret ME, Sjak-Shie EE (2019) Preprocessing pupil size data: Guidelines and code. Behav Res Methods 51:1336–1342.

Kushki A, Drumm E, Mobarak MP, Tanel N, Dupuis A, Chau T, Anagnostou E (2013) Investigating the Autonomic Nervous System Response to Anxiety in Children with Autism Spectrum Disorders. PLOS ONE 8:e59730.

Kuznetsova A, Brockhoff PB, Christensen RHB (2019) lmerTest: Tests in Linear Mixed Effects Models. Available at: https://CRAN.R-project.org/package=lmerTest [Accessed May 25, 2019].

Lawson RP, Mathys C, Rees G (2017) Adults with autism overestimate the volatility of the sensory environment. Nat Neurosci 20:1293–1299.

Lord C, Elsabbagh M, Baird G, Veenstra-Vanderweele J (2018) Autism spectrum disorder. The Lancet 392:508–520.

Martineau J, Hernandez N, Hiebel L, Roché L, Metzger A, Bonnet-Brilhault F (2011) Can pupil size and pupil responses during visual scanning contribute to the diagnosis of autism spectrum disorder in children? J Psychiatr Res 45:1077–1082.

McCloy DR, Larson ED, Lau B, Lee AKC (2016) Temporal alignment of pupillary response with stimulus events via deconvolution. J Acoust Soc Am 139:EL57–EL62.

McGuire RG (1992) Reporting of Objective Color Measurements. HortScience 27:1254–1255.

Michael Waskom et al. (2018) mwaskom/seaborn: v0.9.0 (July 2018). Zenodo. Available at: https://zenodo.org/record/1313201#.XXbuLJNKjGI [Accessed September 9, 2019].

Nuske HJ, Vivanti G, Dissanayake C (2014a) Reactivity to fearful expressions of familiar and unfamiliar people in children with autism: an eye-tracking pupillometry study. J Neurodev Disord 6:14.

Nuske HJ, Vivanti G, Hudry K, Dissanayake C (2014b) Pupillometry reveals reduced unconscious emotional reactivity in autism. Biol Psychol 101:24–35.

Oldfield RC (1971) The assessment and analysis of handedness: The Edinburgh inventory. Neuropsychologia 9:97–113.

Pedregosa F, Varoquaux G, Gramfort A, Michel V, Thirion B, Grisel O, Blondel M, Prettenhofer P, Weiss R, Dubourg V, Vanderplas J, Passos A, Cournapeau D, Brucher M, Perrot M, Duchesnay É (2011) Scikit-learn: Machine Learning in Python. J Mach Learn Res 12:2825–2830.

Pfeffer T, Avramiea A-E, Nolte G, Engel AK, Linkenkaer-Hansen K, Donner TH (2017) Catecholamines Alter the Intrinsic Variability of Cortical Population Activity and Perception. Neuroscience. Available at: http://biorxiv.org/lookup/doi/10.1101/170613 [Accessed June 11, 2019].

Pizzarelli R, Cherubini E (2011) Alterations of GABAergic Signaling in Autism Spectrum Disorders. Neural Plast Available at: https://www.hindawi.com/journals/np/2011/297153/ [Accessed October 26, 2019].

Power JD, Plitt M, Gotts SJ, Kundu P, Voon V, Bandettini PA, Martin A (2018) Ridding fMRI data of motion-related influences: Removal of signals with distinct spatial and physical bases in multiecho data. Proc Natl Acad Sci U S A 115:E2105–E2114.

Remington A, Swettenham J, Campbell R, Coleman M (2009) Selective Attention and Perceptual Load in Autism Spectrum Disorder. Psychol Sci 20:1388–1393.

Revelle W (2019) psych: Procedures for Psychological, Psychometric, and Personality Research. Available at: https://CRAN.R-project.org/package=psych [Accessed May 25, 2019].

Robertson CE, Kravitz DJ, Freyberg J, Baron-Cohen S, Baker CI (2013) Slower Rate of Binocular Rivalry in Autism. J Neurosci 33:16983–16991.

Robertson CE, Ratai E-M, Kanwisher N (2016) Reduced GABAergic Action in the Autistic Brain. Curr Biol CB 26:80–85.

Rosenberg A, Patterson JS, Angelaki DE (2015) A computational perspective on autism. Proc Natl Acad Sci U S A 112:9158–9165.

Samardzic J, Jadzic D, Hencic B, Strac JJ and DS (2018) Introductory Chapter: GABA/Glutamate Balance: A Key for Normal Brain Functioning. GABA Glutamate – New Dev Neurotransmission Res Available at: https://www.intechopen.com/books/gaba-and-glutamate-new-developments-in-neurotransmission-research/introductory-chapter-gaba-glutamate-balance-a-key-for-normal-brain-functioning [Accessed October 26, 2019].

Servan-Schreiber D, Printz H, Cohen JD (1990) A network model of catecholamine effects: gain, signal-to-noise ratio, and behavior. Science 249:892–895.

Sinha P, Kjelgaard MM, Gandhi TK, Tsourides K, Cardinaux AL, Pantazis D, Diamond SP, Held RM (2014) Autism as a disorder of prediction. Proc Natl Acad Sci 111:15220–15225.

Sur M, Rubenstein JLR (2005) Patterning and plasticity of the cerebral cortex. Science 310:805–810.

Urai AE, Braun A, Donner TH (2017) Pupil-linked arousal is driven by decision uncertainty and alters serial choice bias. Nat Commun 8:14637.

Uzunova G, Pallanti S, Hollander E (2016) Excitatory/inhibitory imbalance in autism spectrum disorders: Implications for interventions and therapeutics. World J Biol Psychiatry 17:174–186.

Wagenmakers E-J (2007) A practical solution to the pervasive problems ofp values. Psychon Bull Rev 14:779–804.

Wickham H, François R, Henry L, Müller K, RStudio (2019) dplyr: A Grammar of Data Manipulation. Available at: https://CRAN.R-project.org/package=dplyr [Accessed May 25, 2019].

Williams DL, Goldstein G, Carpenter PA, Minshew NJ (2005) Verbal and Spatial Working Memory in Autism. J Autism Dev Disord 35:747.

Zekveld AA, Koelewijn T, Kramer SE (2018) The Pupil Dilation Response to Auditory Stimuli: Current State of Knowledge. Trends Hear 22 Available at: https://www.ncbi.nlm.nih.gov/pmc/articles/PMC6156203/ [Accessed June 23, 2019].

Zhou S, Yu Y (2018) Synaptic E-I Balance Underlies Efficient Neural Coding. Front Neurosci 12 Available at: https://www.frontiersin.org/articles/10.3389/fnins.2018.00046/full [Accessed October 26, 2019].

